# The selection of reference genome and the search for the origin of SARS-CoV-2

**DOI:** 10.1101/2020.08.10.245290

**Authors:** Yuan Liu, Changhui Yan

## Abstract

The pandemic caused by SARS-CoV-2 has a great impact on the whole world. In a theory of the origin of SARS-CoV-2, pangolins were considered a potential intermediate host. To assemble the coronavirus found in pangolins, SARS-CoV-2 were used a reference genome in most of studies, assuming that pangolins CoV and SARS-CoV-2 are the closest neighbors in the evolution. However, this assumption may not be true. We investigated how the selection of reference genome affect the resulting CoV genome assembly. We explored various representative CoV as reference genome, and found significant differences in the resulting assemblies. The assembly obtained using RaTG13 as reference showed better statistics in total length and N50 than the assembly guided by SARS-CoV-2, indicating that RaTG13 maybe a better reference for assembling CoV in pangolin or other potential intermediate hosts.

## Introduction

Recently, the outbreak of SARS-CoV-2 (COVID-19) has caused an ongoing global pandemic. As of July 21, 2020, the pandemic resulting in a total of 14,562,550 clinical cases and 607,781 deaths all over the world (www.who.int). With an effort of metagenomic RNA deep sequencing, the genome of a SARS-CoV-2 isolate, Wuhan-Hu-1, was published in [11]. The genome showed high nucleotide similarity (89.1%) to a group of SARS-like coronavirus that were identified in bats in China, which indicated the possibility of animal origin.

To identify potential direct or intermediate host of SARS-CoV-2, coronavirus in several animals were studied and compared with SARS-CoV-2. Bat and pangolin were the two mostly investigated species. A CoV in bat, RaTG13, showed 96% full-genome similarity with SARS-CoV-2 [14]. Other virus isolates from bat, ZXC21 and ZC45 also shared 85% of similarity to SARS-CoV-2. These results led to the hypothesis that the progenitor of SARS-CoV-2 originated in Bats and it spilled over to humans using another animal as intermediate host. Many researchers believed that pangolins are a potential intermediate host and they attempted to characterize coronavirus in pangolin. Liu, Chen, and Chen [8] constructed coronavirus contigs using *de novo* assembly method from organ samples of dead Malayan pangolins rescued at the Guangdong Wildlife Rescue Center. Of the 11 collected pangolins, coronavirus was detected in two. Zhang, Wu, and Zhang [13] re-analyzed the RNA-Seq reads from two pangolins carrying coronavirus using reference-guided *de novo* assembly method, with Wuhan-Hu-1 as the reference genome. The resulting draft genome shared 91.02 and 90.55% whole genome similarity with Wuhan-Hu-1 and RaTG13, respectively. Xiao et al. [12] obtained 21 samples from pangolins rescued by Guangdong Customs and conducted reference-guided genome assembly, with Wuhan-Hu-1 as the reference genome. The derived viral genome showed 80 and 98% whole genome sequence identity to SARS-CoV-2 and RaTG13, respectively. Lam et al. [3] collected 43 samples from 18 pangolins from Guangxi Medical University, China, and six samples contained coronavirus sequences. The viral genomes of six samples were *de novo* assembled. They also performed RNA sequencing in five archived pangolins samples from Guangdong, and assembled the genomes using WIV04, another SARS-CoV-2 genome from human, as reference genome. The resultant draft genomes have 85.5% to 92.4% identity to the SARS-CoV-2 genome. They suggested two sub-lineages of coronavirus existed in pangolins. In terms of all coding sites, coronavirus identified in pangolins from Guangdong is more closely related to WIV04 than Bat-CoV RaTG13, whereas coronavirus genome of pangolins in Guangxi showed lower genome similarity to WIV04 than Bat-CoV RaTG13. Liu et al. [8] took samples from three coronavirus positive pangolins rescued in Guangdong and performed deep sequencing. Using *de novo* assembly method, they obtained viral genome that showed 90.32% and 90.24% of whole genome identify to Wuhan-Hu-1and Bat-CoV RaTG13, respectively.

All of the pangolins involved in published coronavirus analysis were from either the Guangdong collection or the Guangxi collection. Pangolins from the Guangdong collection were investigated in most studies [3,8,9,12]. The resulting viral genomes were derived from *de novo* assembly with or without a guided reference and further curated using blast annotation or PCR amplicon sequencing. All studies have showed that CoV in pangolin was highly related to Bat-CoV and SARS-CoV-2. In all of the studies that used reference-guided *de novo* assemblies, a SARS-CoV-2 genome (Wuhan-Hu-1 or WIV04) was chosen as reference [3,12,13]. This choice was based on the assumption that SARS-CoV-2 was the closest neighbor of the Pangolin CoV in the phylogeny tree. However, this assumption may not necessary be true. Therefore, choosing the SARS-CoV-2 genome as reference could inadvertently introduce bias in the genome assembly, leading to inaccurate or incomplete results.

In this study, we assembled the Pangolin CoV genome using several different genomes as reference. We investigated how the reference genome impact the resulting genome and its phylogenetic relationship with others. The results from this study will provide guidance for future studies on how to accurately construct CoV genomes from pangolin or other potential intermediate hosts.

## Material and Methods

### Data selection

Two RNA-seq samples, lung07 and lung08, were downloaded from NCBI SRA under BioProject SRA: PRJNA573298. The two samples were originally published in [8] for viral metagenomics analysis.

### Sequence preparation

Adaptor trimming and quality control were performed on the raw sequence reads using with the Trimmonmatic program (verson 0.39) [1]. To eliminate host contamination, the remaining reads were aligned to the *Manis javanica* genome (SRA: PRJNA256023) using BWA-aln (version 0.7.17) [6] and reads mapped to the host genome were discarded. Reads unmapped to the host reference genome were used to construct genome in the subsequent de novo assembly.

### Genome assembly and sequence analysis

Cleaned reads were used to assemble genome using reference-guided de novo assembly. To investigate how the results were influenced by the choice of reference genome, we explored a few representative virus genomes on the phylogeny tree (Table 1) as the reference genome. These genomes were selected based on previous studies [3,9,11,13]. Once a reference genome was picked, the cleaned reads were aligned to the reference genome using BWA-MEM [5], and the mapped reads were assembled *de novo* using MEGHIT (version 1.1.3) with meta-sensitive mode [4]. The resulting contigs were concatenated into an assembly by aligning them to the reference genome.

**Table 1.**
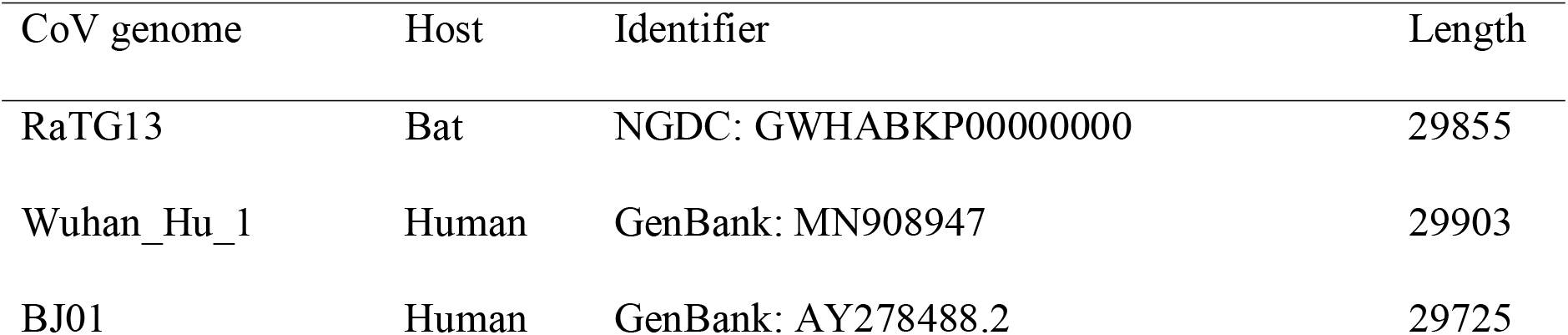
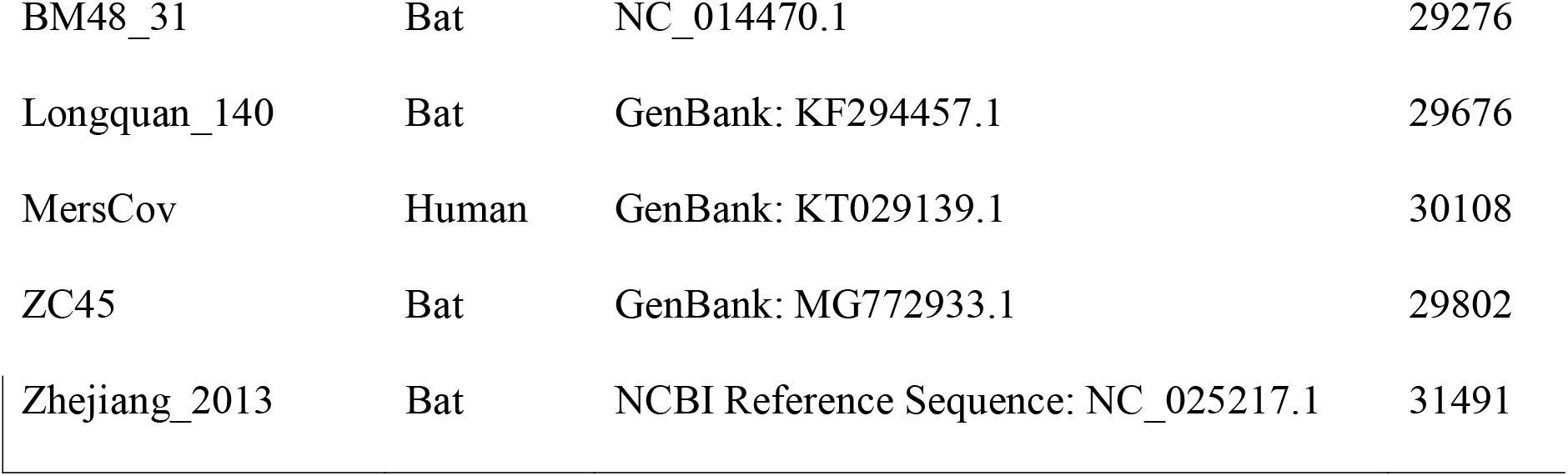
Virus genomes used in the analysis

### Phylogenetic and similarity analyses

Phylogenetic distance analysis was performed using MEGA X (version 10.1.8) [2].The whole genome was used in phylogenetic and distance analysis, and phylogenetic trees were constructed in the best-fit DNA/amino acid substitution mode with 1000 bootstrap replications. The whole genome nucleotide identity analysis was performed in SimPlot 3.5.1 [10].

## Result

A total of eight viral genomes were tested as the reference genome in reference-guided *de novo* assembling. Numbers of mapped reads ranged from 2 to 3,060, and length of the resulting draft assemblies varied from 5,969 to 22,419 bp (Table 2). Two reference genomes, MersCoV and Bat Hp-BetaCoV Zhejiang2013, failed in reads assembling due to the limited number of remaining reads. Less than 1,000 reads were mapped to three reference genomes, BJ01, BM48_31, and Longquan140, resulting in shorter assemblies. A total of 1,061 reads were mapped to ZC45 and subsequently assembled into a 21,819-bp assembly with 67.8% of coverage. RaTG13, which is a bat CoV, had 1,287 reads mapped to it, and the resulting assembly has total length of 21,925 and N50 of 1,428. A total of 3,060 reads were mapped to Wuhan-Hu-1, a human SARS-CoV-2 strain, which was the highest number among all genomes we surveyed. The assembly guided by RaTG13 and Wuhan-Hu-1 showed similar coverage at about eighty percent. However, the resulting assembly had shorter total length (21,819) and N50 (1,195) than those of the assembly that used RaTG13 as reference.

**Table 2.** Summary statistics of assemblies guided by different reference genomes.

| Assembly<br>reference | Reads<br>mapped | Length | N50 | Contig length | Coverage |
| --- | --- | --- | --- | --- | --- |
| BJ01 | 736 | 10100 | 804 | 345-3197 | 0.363 |
| BM48_31 | 368 | 5969 | 518 | 280-1070 | 0.300 |
| Longquan140 | 781 | 12402 | 692 | 280-1683 | 0.421 |
| MersCov | 2 | NA | NA | NA | NA |
| RaTG13 | 1287 | 21925 | 1428 | 322-3464 | 0.790 |
| Wuhan_Hu_1 | 3060 | 21819 | 1195 | 284-3645 | 0.797 |
| ZC45 | 1061 | 21819 | 821 | 277-2436 | 0.678 |
| Zhejiang2013 | 14 | NA | NA | NA | NA |

To understand this seemly contradiction, we investigated the reads coverage and depth on RaTG13 and Wuhan-Hu-1. The results (Figure 1) show that there were an excessive number of reads mapped to distal regions of Wuhan-Hu-1, which could indicate artifacts or contaminations during the sequencing. After removing these tail regions (with > 200X depth), 62 more unique reads were mapped to Wuhan-Hu-1. Figure 2 shows the overlap of between the unique reads mapped to RaTG13 and Wuhan-Hu-1.

**Figure 1.**
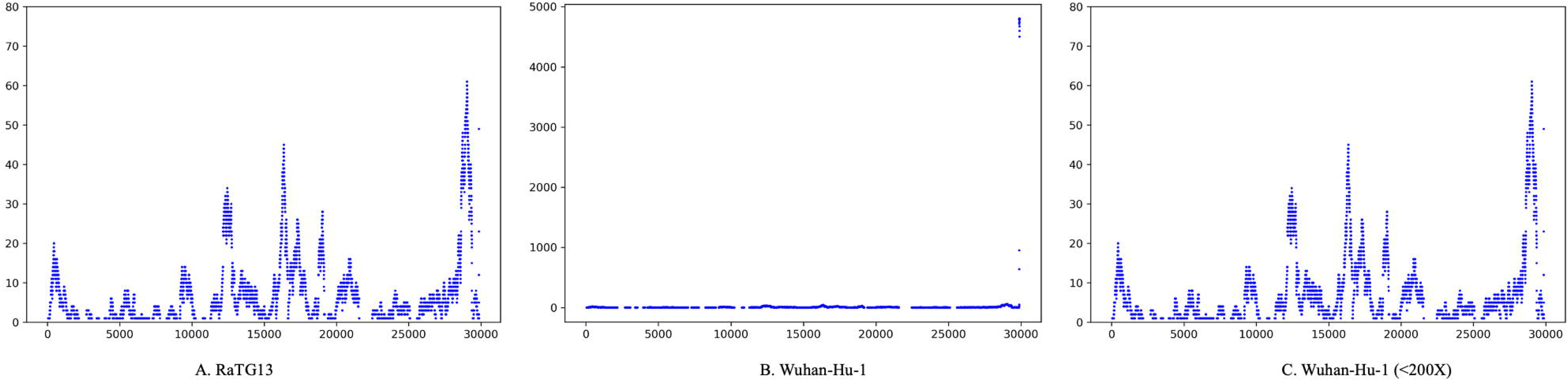
Coverage plot of reads mapped to RaTG13 and Wuhan-Hu-1.

**Figure 2.**
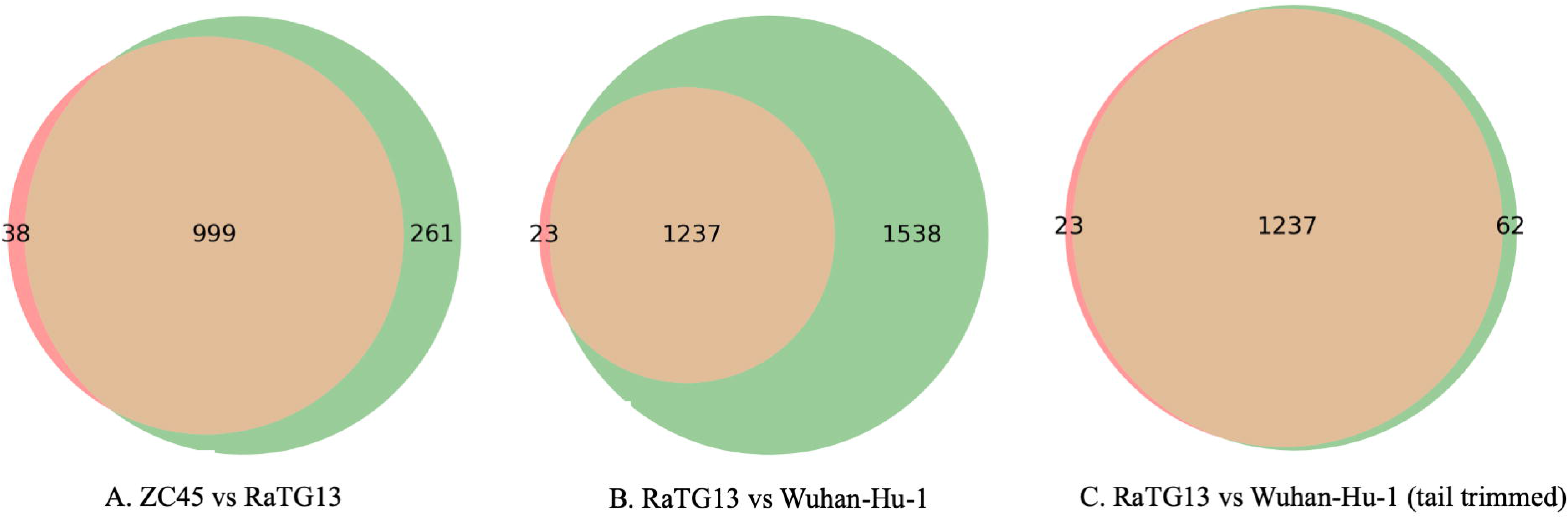
Logical relationships among mapped reads sets.

For further identifying logical relationship of reads mapped to different viral genomes, unique reads were used to construct Venn diagrams. All reads mapped to Italy strain were also mapped to Wuahn-Hu-1 genome (Figure 2). The additional 1516 unique reads were mapped to Wuhan-Hu-1. Most reads were mapped to both RaTG13 and Italy strain, but a total of 45 reads were aligned to either one. Bat-CoV, ZC45 and RaTG13, shared 999 reads in alignment, and RaTG13 genome guided over two hundred more reads into assembly.

The two assemblies that used RaTG13 and Wuhan-Hu-1 as reference genomes respectively, were aligned with the eight reference viral genomes in Table 1 in a multiple alignment. The multiple alignment result was used for similarity analysis and phylogenetic analysis. In terms of the whole genome nucleotide, the RaTG13-guided assembly showed 85.8% and 85.2% identity to RaTG13 and Wuhan-Hu-1, respectively, and the Wuhan-Hu-1-guided assembly showed 88.6 and 88.8% identity to RaTG13 and Wuhan-Hu-1 respectively. The difference between Wuhan-Hu-1-guided assembly and RaTG13-guided assembly was not always consistent with difference between RaTG13 and Wuhan-Hu-1 (Table S1). The differences between RaTG13-guided and Wuhan-Hu-1-guided assemblies in the regions of 18,431-18,601 bp were probably due to references between the reference genomes (Table S1). However, the differences in some regions including 4,761-5,021 bp and 10,121-11,321 bp, were not due to differences in the reference genomes.

We further investigated phylogenetic relationship between the two assemblies and eight reference viral genomes using MEGA X with 1000 Bootstrap tests. The two assemblies positioned between RaTG13 and ZC45 with strong statistical evidence (Figure 3). The SARS-CoV-2 genome, Wuhan-Hu-1, and Bat-CoV, RaTG13, clustered closely together. The two assemblies from this study consistently positioned between this cluster and other CoVs (Figure 4). Among other CoVs, ZC45 is the closest neighbor to the assemblies.

**Figure 3.**
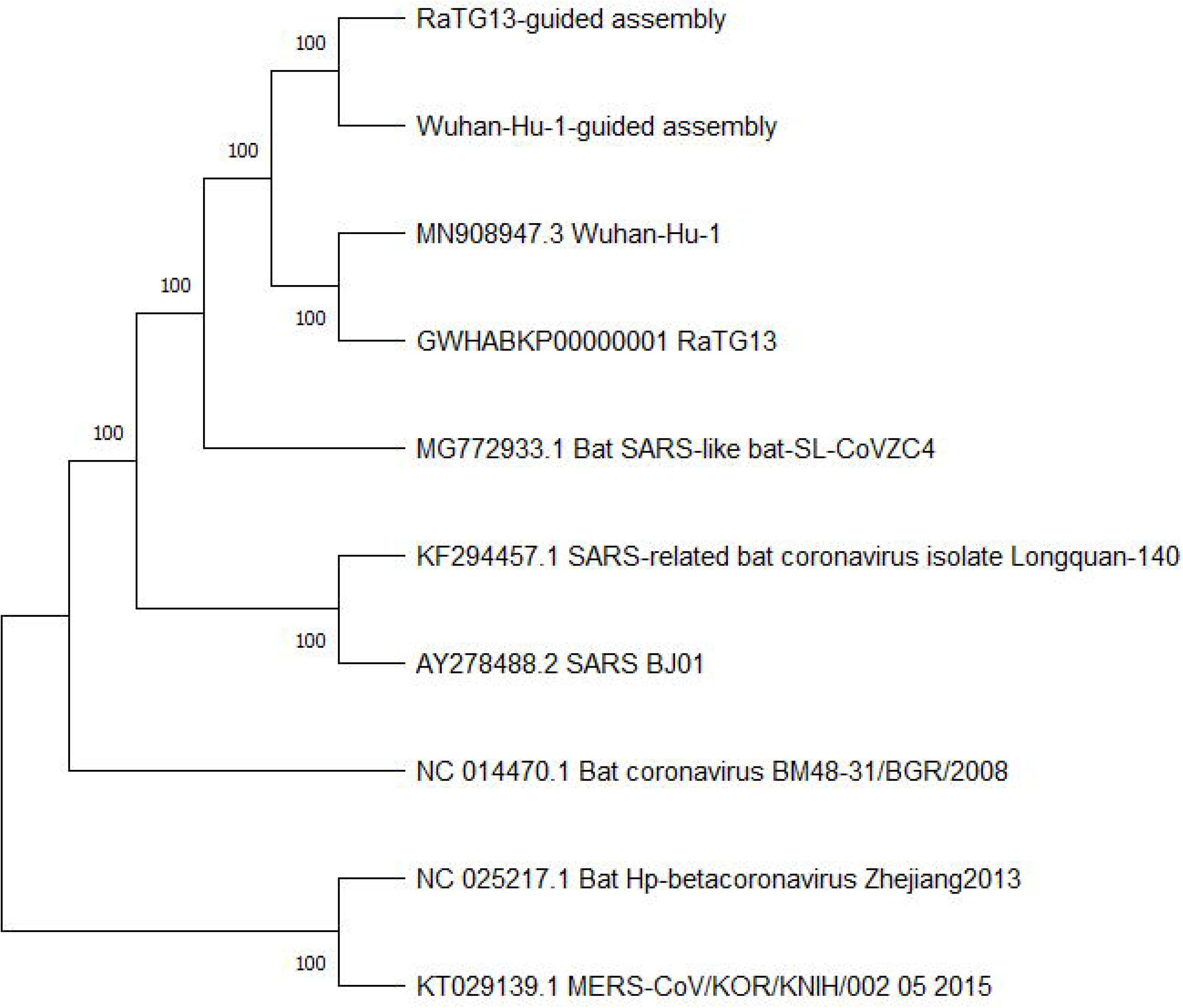
Phylogenetic tree from two assemblies and reference genomes.

## Discussion

Currently, very few samples of coronavirus-positive pangolins have been sequenced. Pangolins-CoV obtained from the Guangdong collection and the Guangxi collection represent two lineages of coronavirus [3]. After outbreak of SARS-CoV, thousands of bat samples were collected and sequenced to identify coronaviruses that bat may carry. We can expect that in near future more pangolin samples will be collected and studied for better understanding of coronavirus in pangolin.

In a reference-guided assembling, the resulting assembly may show bias towards the reference genome [7]. Successful decoding a complete viral genome usually require deep sequencing and further manual curations to fix the gaps. Inaccuracies in assembling the sequencing reads could mislead the subsequent curation step. The whole genome identity between Bat-CoV RaTG13 and SARS-CoV2 Wuhan-Hu-1 is about 96%, which corresponds to a total difference of about 1,500 nucleotides. Our results have shown that observable difference could be found in the resulting assemblies when these genomes were used as reference separately. Particular, when RaTG13 was used as reference, the resulting assembly had a longer total length and higher N50 value than when Wuhan-Hu-1 was used. This points to the possibility that Pangolin-CoV is more closely related to RaTG13 than to Wuhan-Hu-1. Therefore, in order to decode the coronavirus sequence accurately, RaTG13, and possibly other SARS-CoV-2 isolates, should also be considered as reference in future studies of coronavirus in Pangolin or other potential intermediate hosts.

In addition to using one reference genome to guide the assembling, we also attempted to assemble all reads that mapped to either RaTG13 or Wuhan-Hu-1 genomes. The resulting two-genomes-guided assembly has a total length of 22,707 bp, which is slightly longer than that of the RaTG13-guided assembly. However, the N50, 1,388bp, is slightly shorter.

## Abbreviations

CoV: Coronavirus
COVID-19: Coronavirus disease 2019
SARS-CoV-2: Severe acute respiratory syndrome coronavirus
PCR: Polymerase chain reaction

## Conflicts

The authors have no conflict of interest.

---

Table S1. The whole genome nucleotide similarity from RaTG13, Wuhan-Hu-1, and resulting assemblies.

## Notes

### Competing Interest Statement

The authors have declared no competing interest.

## Reference

[1] Bolger, A.M., Lohse, M., Usadel, B. (2014) Trimmomatic: a flexible trimmer for Illumina sequence data. Bioinformatics 30(15), 2114–20, Doi: 10.1093/bioinformatics/btu170.

[2] Kumar, S., Stecher, G., Li, M., Knyaz, C., Tamura, K. (2018) MEGA X: Molecular Evolutionary Genetics Analysis across Computing Platforms. Mol Biol Evol 35(6), 1547–9, Doi: 10.1093/molbev/msy096.

[3] Lam, T.T.-Y., Jia, N., Zhang, Y.-W., Shum, M.H.-H., Jiang, J.-F., Zhu, H.-C., Tong, Y.-G., Shi, Y.-X., Ni, X.-B., Liao, Y.-S., Li, W.-J., Jiang, B.-G., Wei, W., Yuan, T.-T., Zheng, K., Cui, X.-M., Li, J., Pei, G.-Q., Qiang, X., Cheung, W.Y.-M., Li, L.-F., Sun, F.-F., Qin, S., Huang, J.-C., Leung, G.M., Holmes, E.C., Hu, Y.-L., Guan, Y., Cao, W.-C. (2020) Identifying SARS-CoV-2-related coronaviruses in Malayan pangolins. Nature, 1–4, Doi: 10.1038/s41586-020-2169-0.

[4] Li, D., Liu, C.-M., Luo, R., Sadakane, K., Lam, T.-W. (2015) MEGAHIT: an ultra-fast single-node solution for large and complex metagenomics assembly via succinct de Bruijn graph. Bioinformatics 31(10), 1674–6, Doi: 10.1093/bioinformatics/btv033.

[5] Li, H. (2013) Aligning sequence reads, clone sequences and assembly contigs with BWA-MEM. ArXiv:1303.3997 [q-Bio].

[6] Li, H., Durbin, R. (2009) Fast and accurate short read alignment with Burrows-Wheeler transform. Bioinformatics 25(14), 1754–60, Doi: 10.1093/bioinformatics/btp324.

[7] Lischer, H.E.L., Shimizu, K.K. (2017) Reference-guided de novo assembly approach improves genome reconstruction for related species. BMC Bioinformatics 18(1), 474, Doi: 10.1186/s12859-017-1911-6.

[8] Liu, P., Chen, W., Chen, J.-P. (2019) Viral Metagenomics Revealed Sendai Virus and Coronavirus Infection of Malayan Pangolins (Manis javanica). Viruses 11(11), 979, Doi: 10.3390/v11110979.

[9] Liu, P., Jiang, J.-Z., Wan, X.-F., Hua, Y., Li, L., Zhou, J., Wang, X., Hou, F., Chen, J., Zou, J., Chen, J. (2020) Are pangolins the intermediate host of the 2019 novel coronavirus (SARS-CoV-2)? PLOS Pathogens 16(5), e1008421, Doi: 10.1371/journal.ppat.1008421.

[10] Lole, K.S., Bollinger, R.C., Paranjape, R.S., Gadkari, D., Kulkarni, S.S., Novak, N.G., Ingersoll, R., Sheppard, H.W., Ray, S.C. (1999) Full-Length Human Immunodeficiency Virus Type 1 Genomes from Subtype C-Infected Seroconverters in India, with Evidence of Intersubtype Recombination. Journal of Virology 73(1), 152–60, Doi: 10.1128/JVI.73.1.152-160.1999.

[11] Wu, F., Zhao, S., Yu, B., Chen, Y.-M., Wang, W., Song, Z.-G., Hu, Y., Tao, Z.-W., Tian, J.-H., Pei, Y.-Y., Yuan, M.-L., Zhang, Y.-L., Dai, F.-H., Liu, Y., Wang, Q.-M., Zheng, J.-J., Xu, L., Holmes, E.C., Zhang, Y.-Z. (2020) A new coronavirus associated with human respiratory disease in China. Nature 579(7798), 265–9, Doi: 10.1038/s41586-020-2008-3.

[12] Xiao, K., Zhai, J., Feng, Y., Zhou, N., Zhang, X., Zou, J.-J., Li, N., Guo, Y., Li, X., Shen, X., Zhang, Z., Shu, F., Huang, W., Li, Y., Zhang, Z., Chen, R.-A., Wu, Y.-J., Peng, S.-M., Huang, M., Xie, W.-J., Cai, Q.-H., Hou, F.-H., Chen, W., Xiao, L., Shen, Y. (2020) Isolation of SARS-CoV-2-related coronavirus from Malayan pangolins. Nature, Doi: 10.1038/s41586-020-2313-x.

[13] Zhang, T., Wu, Q., Zhang, Z. (2020) Probable Pangolin Origin of SARS-CoV-2 Associated with the COVID-19 Outbreak. Current Biology 30(7), 1346–1351.e2, Doi: 10.1016/j.cub.2020.03.022.

[14] Zhou, P., Yang, X.-L., Wang, X.-G., Hu, B., Zhang, L., Zhang, W., Si, H.-R., Zhu, Y., Li, B., Huang, C.-L., Chen, H.-D., Chen, J., Luo, Y., Guo, H., Jiang, R.-D., Liu, M.-Q., Chen, Y., Shen, X.-R., Wang, X., Zheng, X.-S., Zhao, K., Chen, Q.-J., Deng, F., Liu, L.-L., Yan, B., Zhan, F.-X., Wang, Y.-Y., Xiao, G.-F., Shi, Z.-L. (2020) A pneumonia outbreak associated with a new coronavirus of probable bat origin. Nature 579(7798), 270–3, Doi: 10.1038/s41586-020-2012-7.

